# A Pilot Survey of Authors’ Experiences with Poor Peer Review Practices

**DOI:** 10.1101/2022.12.20.521261

**Authors:** Kyle McCloskey, Jon F. Merz

## Abstract

**Objectives:** To develop a typology of poor peer review practices (PPRP) and assess researchers’ experiences with PPRP.

**Design:** Exploratory analysis of cross-sectional internet-based survey.

**Participants:** We solicited 500 researchers funded by the NIH extramural grants in 2018 by direct email and 600 bioethicists on a bioethics discussion forum (mcw.bioethics). 112 respondents (~10%) completed the survey.

**Primary and Secondary Outcomes Measures:** The total number of reported PPRP and a five-point scale to assess participants’ views about the effect of PPRP on their ability to disseminate their research.

**Results:** The mean number of PPRP experienced per author was 12.5 of 28 (44.6%; range 0–27; 95% CI = 11.2–13.8), with fourteen PPRP experienced by 50% or more of the sample. The number of reported PPRP increased with age (P = 0.01) and total number of published peer-reviewed manuscripts (P = 0.02). Authors belonging to underrepresented groups reported more PPRP compared to represented groups (P = 0.05). Most authors viewed the peer review process favorably, with 67% (74/111) of authors responding “sometimes” or “often” to having received insightful peer reviews that improved the quality of their final papers. However, a total of 57% (63/111) of respondents admitted to previously abandoning a manuscript after receiving what they perceived to be unfair peer reviews.

**Conclusions:** This study introduces a practical list of PPRP and a framework for a typology of PPRP, which could serve as an educational tool for editors and reviewers and further our understanding of poor peer review practices. Future researchers will expand authors’ experiences with constructive or helpful peer review practices.

**STRENGTHS AND LIMITATIONS OF THIS STUDY:**

- The sample consisted primarily of experienced researchers from diverse fields, which aided in capturing a wide variety of poor peer review examples.
- This survey included a core set of 28 poor peer review practices and allowed respondents to add other practices they had experienced, which helped to generate an extensive list of poor peer review practices.
- The generalizability of the prevalence of poor review types and the degree of negative impact on authors should be interpreted with caution due to the low response rate and the potential for response bias.

## 1. INTRODUCTION

Peer review is a fundamental tool of the editorial process that helps maintain scientific integrity. Since its introduction in 1731 by the Royal Society of Edinburg, it has become the “gold standard” for evaluating scholarly work by calling on independent reviewers to assess a study’s validity, quality, and originality.^1^ This process is intended to prevent the publication of inaccurate or poor-quality research while providing expert feedback to authors to aid them in producing higher-quality work.^1–3^

Over the past few decades, however, a growing body of literature has begun to confirm the various limitations of the peer review process, from biased or inexperienced reviewers to low inter-reliability between reviews.^4–9^ The little guidance offered to peer reviewers on what constitutes a “good” or “poor” peer review may also partly explain these shortcomings. While more educational resources have become available for reviewers, it is still axiomatic that reviewers occasionally provide ambiguous critiques or cross the line, making comments that strike authors as bothersome or worse.^9–13^ For authors, these limitations can make peer review a frustrating and mysterious process that deters them from disseminating their work. It is perhaps axiomatic that authors set aside reviews for a few days after receipt to calm the agita and avoid quick and impassioned responses.

To begin to explore these issues, we performed a pilot cross-sectional survey study to assess the types of poor peer review practices (PPRP) encountered by researchers, to develop a typology of poor peer review practices, and to begin to assess the impact of poor peer review practices on authors’ perceived abilities to disseminate their research. A pilot study was performed to test the survey instrument before a larger-scale study. This knowledge could potentially serve as an educational tool to elucidate issues hindering the peer review process from its goal of promoting the quality and integrity of the sciences.

## 2. METHODS

Data is reported according to the Consensus-Based Checklist for Reporting of Survey Studies (CROSS).^14^ The institutional review board at the University of Pennsylvania deemed this study exempt.

### 2.1 Study participants

An invitation to complete the survey was emailed to a random sample of 500 researchers funded by the National Institute of Health’s (NIH) extramural grants in 2018, with replacement of emails returned undeliverable.^15^ An additional invitation was posted to a bioethics discussion forum with roughly 500 members to increase the diversity of researcher disciplines in the sample (Art Derse, personal communication).

### 2.2 Survey design

An anonymous three-section, 51-question survey was designed using Qualtrics (Seattle, Washington, USA). The first section assessed the prevalence of 28 unique PPRP examples through yes-no questions (e.g., “Have you experienced the following examples of poor peer review practices?” – e.g., “*Ad hominem* attack,” “Unbalanced negative review,” “Unstructured review”). The 28 unique PPRP examples were compiled based on the experience of one of the authors (JFM), a literature review, and a scan of the Facebook user group Reviewer2mustbestopped. Participants were also asked open-ended questions about whether they experienced poor peer reviews not listed in the survey to capture any additional types of PPRP missed in the author-generated list.

The second section utilized Likert scale questions to assess the effect of PPRP on participants’ ability to conduct and disseminate their research. Participants were asked to rate six statements on a five-point scale (e.g., “How would you rate the following statement: ‘I have considered leaving academia after receiving unfair review.’? – “never,” “rarely,” “sometimes,” “often,” “all the time”). The last section of the survey asked about demographic information (age, gender, race, language, field of study, total number of peer-reviewed publications, category of types of peer-reviewed publication) through multiple-choice questions. For data on underrepresented demographics, participants were asked the yes-no question: “Do you consider yourself a part of an underrepresented demographic in your field of work?”.

The presentation of questions was randomized for each section except for demographic questions. No incentives or prizes were offered for completing the survey, and consent was required from each subject before entering the survey. Each respondent was restricted to only one submission. The final instrument was pre-tested using two expert reviewers, which helped identify missing topics in the survey and determine content and response process validity. Three mailings were performed at one-week intervals throughout April 2022, and data was collected for analysis at the end of April 2022. The survey instrument is provided in the supplementary material.

### 2.3 Statical analyses and categorization of poor peer review practices

Respondents reported PPRP experiences were summarized and compared across various demographic characteristics using an exploratory nonparametric Wilcoxon rank-sum (Mann-Whitney) test and a Cuzick extension of Wilcoxon rank-sum test for ordered groups (Stata 12.1, StataCorp, © 2014). Two-sided P < .05 was considered significant, and consistent with the exploratory nature of this analysis, P values were unadjusted for multiple comparisons. In addition to statical analyses, the authors categorized the examples of PPRP reported by participants according to similar themes to help process and understand the diversity of PPRP examples.

## 3. RESULTS

### 3.1 Respondent characteristics

A total of 112 researchers completed the surveys, roughly 10% of those solicited (**Table**). The respondents were predominantly male (54%), held a Ph.D. (78%), > 50 years old (66%), identified as white (87%), published more than 50 peer-reviewed papers in their career (64%), were trained in the humanities or social sciences (54%) and conducted primarily empirical research (61%).

**Table.** Univariate exploratory analysis of total poor peer review practices (PPRP)

| Factor (N) | PPRP:Mean (SD) | Z score | P value |
| --- | --- | --- | --- |
| **Age** |  |  |  |
| 20-29 (9) | 15.4 (5.9) | -2.51 | 0.01<sup>b</sup> |
| 30-39 (29) | 14.7 (7.0) |  |  |
| 40-49 (33) | 12.0 (7.0) |  |  |
| 50-59 (40) | 10.8 (6.1) |  |  |
| **Gender** |  |  |  |
| Female (50) | 11.5 (6.9) | 1.39 | 0.16<sup>a</sup> |
| Male (59) | 13.3 (6.6) |  |  |
| **Race** |  |  |  |
| White (92) | 12.0 (7.0) | 1.62 | 0.10<sup>a</sup> |
| Non-white (14) | 15.0 (4.9) |  |  |
| **Underrepresented group** |  |  |  |
| No (84) | 11.8 (7.1) | 1.97 | 0.05<sup>a</sup> |
| Yes (22) | 14.8 (5.2) |  |  |
| **Career peer-reviewed publications** |  |  |  |
| ≤ 10 | 10.8 (5.8) | 2.37 | 0.02<sup>b</sup> |
| 11-50 | 11.1 (6.8) |  |  |
| 51-100 | 11.9 (6.9) |  |  |
| >100 | 14.7 (6.6) |  |  |
| **Have received positive, helpful peer reviews:** |  |  |  |
| Never (3) | 16.0 (5.0) | -2.53 | 0.01<sup>b</sup> |
| Rarely (34) | 14.0 (7.3) |  |  |
| Sometimes (65) | 12.3 (6.4) |  |  |
| Often (9) | 7.7 (5.0) |  |  |

a = Nonparametric Wilcoxon rank-sum (Mann-Whitney) test; b = Cuzick extension of Wilcoxon rank-sum test for ordered groups.

### 3.2 Interactions between the number of poor peer reviews practices and other variables

The mean number of PPRPs experienced per author was 12.5 of 28 (44.6%; range 0–27; 95% CI = 11.2–13.8), with fourteen PPRPs experienced by 50% or more of the sample (**Figure 1**). As shown in the exploratory results presented in **Table,** reported PPRP increased with age (P = 0.01), the total number of published peer-reviewed manuscripts (P = 0.02), in self-identified underrepresented groups (P = 0.05), and those who received fewer “helpful/positive” reviews (P = 0.01). No statistical differences were found in the number of reported PPRPs between race and gender.

**Figure 1.**
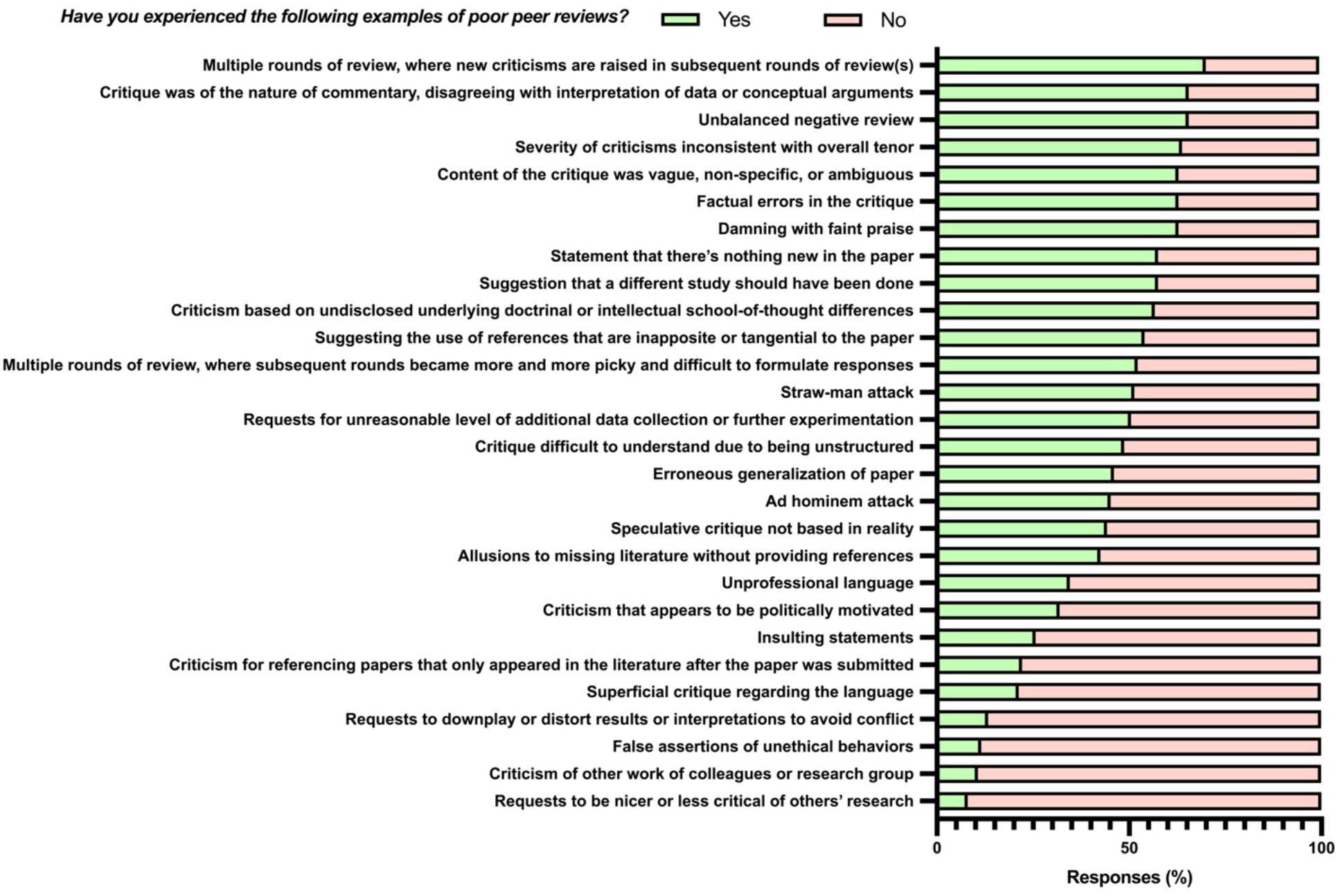
Prevalence of poor peer review practices assessed through yes-no questions.

### 3.3 Impact of poor peer review practices on authors

Most authors viewed the peer review process favorably, with 67% (74/111) of authors responding “sometimes” or “often” to having received insightful peer reviews that improved the quality of their final papers. However, a total of 57% (63/111) of respondents admitted to previously abandoning a manuscript after receiving unfair peer reviews. Moreover, 72% (81/112) of respondents admitted feeling discouraged after receiving unfair peer reviews, though 33% (37/112) said they are rarely discouraged. As shown in **Table**, the total number of PPRP reported by respondents increased with the number of reported adverse consequences (p=0.001). A further examination of PPRP on authors can be found in **Figure 2**.

**Figure 2.**
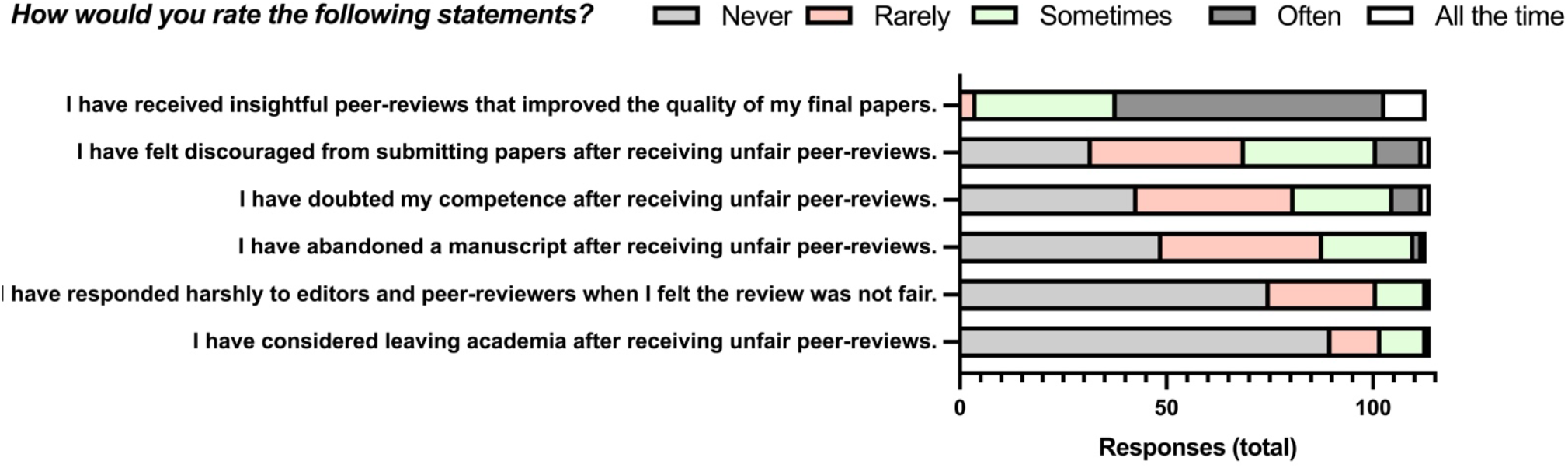
Authors’ experiences with poor peer review practices assessed through 5-point Likert-style questions.

### 3.4 A typology of poor peer review practices

A typology of PPRP was developed to understand the extensive range of PPRP experienced by respondents. Five notable themes in reported PPRP examples were summarized:

1. *Ill-natured Reviews* – Peer reviews that have an unprofessional disposition and appear to be an explicit or implicit attack on the author’s work or intellectual schools of thought, such as *ad hominem* attacks or insulting statements.
2. *Erroneous Reviews* – Peer reviews that contain factual errors, such as false assertions of unethical behavior, misreading of findings, or erroneous generalizations of the paper.
3. *Unreasonable Requests* – Peer reviews with unnecessary or unreasonable requests, such as asking for extensive additional data collection or experimentation, suggestions that a different study should have been done, or requests to cite irrelevant (and presumably a reviewer’s) work.
4. *Inconsequential or Incoherent Reviews* – Peer reviews that are overly particular and do not impact the overall argument of the work or are incoherent due to being unstructured, poorly written, non-specific, or vague.
5. *Editorial or Process Issues* – Issues internal to the editorial process and negatively impact authors’ ability to disseminate their work or result, such as an insufficient number of peer reviewers, breaks of anonymity, or multiple rounds of peer review where new criticisms are raised on subsequent rounds.

## 4. DISCUSSION

To our knowledge, this is the first study to systematically analyze the types of poor peer review practices and develop a typology of PPRP. These findings suggest that researchers may encounter an extensive range of PPRP. The sample consisted primarily of experienced researchers from diverse fields, which likely aided in capturing a wide variety of poor peer review examples. Indeed, respondents who published more peer-reviewed manuscripts also reported experiencing more PPRP. The list and typology of poor peer review practices introduced in this study could serve as an educational tool for editors and reviewers and further our understanding of poor peer review practices.

The most commonly reported practice was “Multiple rounds of peer review where new criticisms are raised in subsequent rounds” and is classified as an *Editorial or Process Issue.*While some editors may see this as a normal function of the peer review process,^1,2^ authors may view it as a cumbersome issue that delays the dissemination of their work. New criticisms raised on subsequent rounds of revisions could potentially reflect the low reliability between reviewers and the inherent arbitrariness of peer review; however, there are many reasons why new criticisms are raised throughout the review process, with some justified and others unnecessary. Editors should thus consider such issues on a case-by-case basis. While *Ill-natured, Erroneous,*and *Incoherent Reviews* are unequivocally poor peer review practices that do not warrant further discussion, there is a need for further research on what constitutes a PPRP for *Editorial or Process Issues* that are more ambiguous.

This study also investigated the impact of the peer review process on authors’ ability to disseminate their research. Unsurprisingly, most respondents benefited from the peer review process, with 58% of authors responding “often” to receiving insightful peer reviews that improved the quality of their final papers. However, this finding was tempered by 57% of authors who admitted to abandoning a manuscript after receiving unfair peer reviews. The findings also suggest that authors who self-identify as underrepresented minorities in their respective fields are more likely to report experiencing PPRP than represented groups. Previous research found similar findings, with unprofessional peer reviews disproportionately harming underrepresented groups in STEM.^8^ These findings confirm that the peer review process is indeed a powerful tool of the editorial process that can improve a researcher’s work; however, it may stymie researchers’ ability to disseminate their research when misused.

The generalizability of the prevalence of poor review types and the degree of negative impact on authors should be interpreted with caution due to the low response rate and the potential for response bias, given the nature of survey studies. However, the low response rate is less germane to the primary aim of generating a list and typology of poor peer review practices. Another limitation is that this study focused on poor peer review practices and excluded any in-depth exploration of constructive or helpful peer review practices. Further research is needed to establish greater external validity of the prevalence of the types of poor peer review practices generated in this study and their effect on researchers’ ability to disseminate their research. Furthermore, while this pilot study confirmed the survey instrument’s feasibility and efficacy, we plan to expand the survey to explore constructive or helpful peer review practices and examine their effects on early-career researchers.

## 5. CONCLUSION

This study introduces a practical list of poor peer review practices and a framework for a typology of poor peer review practices, which could serve as an educational tool for editors and reviewers and further our understanding of poor peer review practices. These findings also suggest that researchers may encounter an extensive range of poor peer reviews of various types. Further research is warranted on *Editorial and Process Issues* that are more ambiguous and, consequently, difficult to assess whether their function promotes or hinders the peer review process. Future researchers should also consider investigating authors’ experiences with constructive or helpful peer review practices.

## Supporting information

Survey Instrument

## AUTHOR CONTRIBUTIONS

**Kyle McCloskey:** Writing - original draft, Writing - review & editing, Conceptualization, Visualization. **Jon F. Merz**: Writing - original draft, Writing - review & editing, Conceptualization, Project administration, Supervision, Resources.

## ACKNOWLEDGEMENTS

The authors thank scientist and bioethicist respondents for completing the survey, Jim Coyne, and anonymous reviewers for comments. An abstract presenting these findings were posted at the Ninth International Congress on Peer Review and Scientific Publication, Chicago, IL, USA, September 8-10, 2022. Responsibility for the work is solely that of the authors.

