## Supplementary material for "A Pilot Survey of Authors’ Experiences with Poor Peer Review Practices": Survey Instrument

Start of Block: Block 1

We would like you to take part in a brief survey study about peer review. We’re trying to get a sense of the frequency and ramifications of poor quality, obnoxious, or upsetting peer review practices. This should take no more than **10 minutes** of your time. 
 
This survey is **anonymous**. You received our solicitation because you: 1) are an NIH-funded investigator; or 2) are a member of a professional society or internet discussion forum that has allowed us to distribute our request. You may receive more than one request; if you do, please only reply once. 
 
This survey has been determined to be exempt by the University of Pennsylvania Institutional Review Board. If you have any questions or would like a copy of the results of this project, please contact us. 


Thank you very much! We appreciate your time and effort.   
 
Jon Merz:; 267-254-6470  
Kyle McCloskey: 
 
**Click "I agree" to take part.**

- I agree (1)
- I do not agree (2)

Skip To: End of Survey If We would like you to take part in a brief survey study about peer review. We’re trying to get a s... = I do not agree

End of Block: Block 1

Start of Block: Block 2

Have you encountered the following examples of ill-natured reviews?


*Note: we are focused on peer-review of authored journal articles or longer manuscripts, and not grant applications. We do not expect you to pull out old peer reviews to answer the following questions, so please answer all questions from recall.*

Damning with faint praise (e.g., not finding fault, but giving the paper a big “so what?”).

- No (1)
- Yes (2)

Ad hominem attack (e.g., suggestion that the author(s) do not know what they are talking about).

- No (1)
- Yes (2)

Insulting statements (e.g., derogatory statements about the author(s), institutions, the research, etc.).

- No (1)
- Yes (2)

Unprofessional language (e.g., abusive, obnoxious, disrespectful, hurtful, sarcastic comments).

- No (1)
- Yes (2)

Severity of criticisms inconsistent with overall tenor/recommendations.

- No (1)
- Yes (2)

Unbalanced negative review (i.e., the review only focused on criticism without highlighting what was done well).

- No (1)
- Yes (2)

Criticism of other (perhaps unrelated) work of colleagues or research group.

- No (1)
- Yes (2)

Criticism that appears to be politically motivated (e.g., unpopular findings; protection of a ‘sacred cow’).

- No (1)
- Yes (2)

Criticism based on undisclosed underlying doctrinal or intellectual school-of-thought differences.

- No (1)
- Yes (2)

End of Block: Block 2

Start of Block: Block 3

Have you encountered the following examples of erroneous reviews?

False assertions of unethical behaviors (e.g., double publication, plagiarism, violation of human subjects or animal protections, etc.).

- No (1)
- Yes (2)

Factual errors in the critique (i.e., making critical statements that are wrong, with no supporting references).

- No (1)
- Yes (2)

Erroneous generalization of paper (i.e., not interpreting the paper’s findings as a whole and unfairly narrowing their critique to a section).

- No (1)
- Yes (2)

Straw-man attack (i.e., misinterpreting the paper and attacking the misinterpretation).

- No (1)
- Yes (2)

Criticism for not referencing papers that only appeared in the literature after the paper was submitted.

- No (1)
- Yes (2)

Critique was of the nature of commentary, disagreeing with interpretation of data or conceptual arguments.

- No (1)
- Yes (2)

Suggesting the use of references that are inapposite or tangential to the paper (perhaps the reviewer’s own works).

- No (1)
- Yes (2)

Speculative critique not based in reality (e.g., what-about-isms).

- No (1)
- Yes (2)

End of Block: Block 3

Start of Block: Block 4

Have you encountered the following examples of unreasonable requests in a peer-review?

Suggestion that a different study should have been done, not the one that was done.

- No (1)
- Yes (2)

Requests for unreasonable level of additional data collection or further experimentation.

- No (1)
- Yes (2)

Multiple rounds of review, where subsequent rounds became more and more picky and difficult to formulate responses.

- No (1)
- Yes (2)

Multiple rounds of review, where new criticisms are raised in subsequent rounds of review(s).

- No (1)
- Yes (2)

Requests to downplay or distort results or interpretations to avoid conflict.

- No (1)
- Yes (2)

Requests to be nicer or less critical of others’ research.

- No (1)
- Yes (2)

End of Block: Block 4

Start of Block: Block 5

Have you encountered the following examples of incoherent or **inconsequential reviews?**

Allusions to missing literature without providing references.

- No (1)
- Yes (2)

Content of the critique was vague, non-specific, or ambiguous.

- No (1)
- Yes (6)

Statement that there’s nothing new in the paper.

- No (1)
- Yes (2)

Critique difficult to understand due to being unstructured.

- No (1)
- Yes (2)

Superficial critique regarding the language (e.g., lack of academic terminology).

- No (1)
- Yes (2)

End of Block: Block 5

Start of Block: Block 6

**You've finished half of the survey already!**

Have you received any other review(s) that upset you or struck you as unfair that was not previously mentioned?

- No (4)
- Yes (5)

Display This Question:

If Have you received any other review(s) that upset you or struck you as unfair that was not previou... = Yes

Please describe the unfair review(s). *Note: we do not expect you to pull out old reviews to answer this question, please summarize the review from recall.*

________________________________________________________________

Have you ever appealed rejection of an article because the peer review(s) failed in some way?

- No (4)
- Yes (5)

Display This Question:

If Have you ever appealed rejection of an article because the peer review(s) failed in some way?  = Yes

Please share details about your appealed rejection (optional).

________________________________________________________________

Considering the examples of poor peer-review practices mentioned in this survey, do you think that you have engaged in any of these *as a reviewer*?

- I never wrote a review (1)
- Never (2)
- I've tried not to, but there have been times (10)
- Yes, I don't think reviewers should hold back (5)
- Prefer not to say (9)

End of Block: Block 6

Start of Block: Block 7

**How would you rate the following statements with a scale from never—rarely —sometimes—often—all the time?**

I have doubted my competence after receiving unfair peer-reviews.

- Never (1)
- Rarely (2)
- Sometimes (3)
- Often (4)
- All the time (5)

I have felt discouraged from submitting papers after receiving unfair peer-reviews.

- Never (1)
- Rarely (2)
- Sometimes (3)
- Often (4)
- All the time (5)

I have abandoned a manuscript after receiving unfair peer-reviews.

- Never (1)
- Rarely (2)
- Sometimes (3)
- Often (4)
- All the time (5)

I have considered leaving academia after receiving unfair peer-reviews.

- Never (1)
- Rarely (2)
- Sometimes (3)
- Often (4)
- All the time (5)

I have responded harshly to editors and peer-reviewers when I felt the review was not fair.

- Never (1)
- Rarely (2)
- Sometimes (3)
- Often (4)
- All the time (5)

I have received insightful peer-reviews that improved the quality of my final papers.

- Never (1)
- Rarely (2)
- Sometimes (3)
- Often (4)
- All the time (5)

End of Block: Block 7

Start of Block: Block 8

**This survey is almost complete. Thank you for your participation thus far.**

How many peer reviews of journal articles in your field have you performed in your career?

- 0 (6)
- 1 - 10 (2)
- 11 - 50 (3)
- 51 - 100 (4)
- > 100 (5)

About how many journal peer reviews have you performed in the last year?

- 0 (1)
- 1 - 6 (2)
- > 7 (3)

How many peer-reviewed articles have you published in your career?

- 1 - 10 (1)
- 11 - 50 (2)
- 51 - 100 (3)
- > 100 (4)

Select the field(s) that best describe your area of research.

- Applied sciences (e.g., medicine, engineering) (4)
- Natural sciences (e.g., biology, chemistry, physics, and earth science) (1)
- Social sciences (e.g., psychology, sociology, economics) (2)
- Humanities (e.g., philosophy, history, law) (5)
- Formal sciences (e.g., mathematics, theoretical computer science) (3)

About what percentage of your peer-reviewed publications involves the following intellectual activities.

Conceptual or theoretical work (e.g., philosophy, law, religious studies, other) : _______ (1)

Editorials or commentary : _______ (2)

Reviews (e.g., systematic reviews) : _______ (3)

Empirical studies (involving primary collection of data) : _______ (4)

Other : _______ (5)

Total : ________

Have you served as an editor for a journal?

- No (1)
- Yes (2)

End of Block: Block 8

Start of Block: Block 9

**A few background questions to finish!**

How old are you?

- 20 - 29 (1)
- 30 - 39 (2)
- 40 - 49 (3)
- 50 - 59 (4)
- > 60 (5)

How do you describe yourself?

- Male (1)
- Female (2)
- Non-binary / third gender (3)
- Prefer not to say (4)

Is English your primary language?

- Yes (1)
- No (2)
- Prefer not to say (3)

Choose one or more races and/or ethnicities that you identify with:

- White (1)
- Black or African American (2)
- American Indian or Alaska Native (3)
- Asian (4)
- Native Hawaiian or Pacific Islander (5)
- Hispanic or Latinx (6)
- Prefer not to say (7)
- Other (8) __________________________________________________

Do you consider yourself a member of an underrepresented demographic in your field of research?

- Yes (1)
- No (2)
- Prefer not to say (3)

What is your highest level of educational attainment (please check all that apply)?

- BA/BS (1)
- MA/MS (2)
- MD, DO, DVM, or equivalent (3)
- PhD, DPH, or equivalent (4)
- JD/LLB/LLM/LLD (5)
- Other (6) __________________________________________________

End of Block: Block 9
